# Analysis of Mutations in Precision Oncology using The Automated, Accurate, and User-Friendly Web Tool PredictONCO

**DOI:** 10.1101/2024.06.08.598056

**Authors:** Rayyan Tariq Khan, Petra Pokorna, Jan Stourac, Simeon Borko, Adam Dobias, Joan Planas-Iglesias, Stanislav Mazurenko, Ihor Arefiev, Gaspar Pinto, Veronika Szotkowska, Jaroslav Sterba, Jiri Damborsky, Ondrej Slaby, David Bednar

## Abstract

Next-generation sequencing technology has created many new opportunities for clinical diagnostics, but it faces the challenge of functional annotation of identified mutations. Various algorithms have been developed to predict the impact of missense variants that influence oncogenic drivers. However, computational pipelines that handle biological data must integrate multiple software tools, which can add complexity and hinder non-specialist users from accessing the pipeline. Here, we have developed an online user-friendly web server tool PredictONCO that is fully automated and has a low barrier to access. The tool models the structure of the mutant protein in the first step. Next, it calculates the protein stability change, pocket level information, evolutionary conservation, and changes in ionisation of catalytic amino acid residues, and uses them as the features in the machine-learning predictor. The XGBoost-based predictor was validated on an independent subset of held-out data, demonstrating areas under the receiver operating characteristic curve (ROC) of 0.95 and 0.94, and the average precision from the precision-recall curve 0.98 and 0.94 for structure-based and sequence-based predictions, respectively. Finally, PredictONCO calculates the docking results of small molecules approved by regulatory authorities. We demonstrate the applicability of the tool by presenting its usage for variants in two cancer-associated proteins, cellular tumour antigen p53 and fibroblast growth factor receptor FGFR1. Our free web tool will assist with the interpretation of data from next-generation sequencing and navigate treatment strategies in clinical oncology: https://loschmidt.chemi.muni.cz/predictonco/.

## Introduction

In the last two decades, we have witnessed substantial technological advancements in human genomics attributed mainly to the implementation of next-generation sequencing (NGS). With its ability to simultaneously analyse a large amount of genetic information, increasing availability, and decreasing costs, NGS has already been adopted by multiple academic and clinical laboratories and is getting to the forefront of medical diagnostics. This considerable progress and the resulting impact on clinical management is especially apparent in oncology, where comprehensive genomic profiling brings valuable information on the presence of acquired somatic alterations that can be utilised for therapeutic planning within the paradigm of precision medicine [1], with further augmentation of predictive capabilities by artificial intelligence [2].

Several knowledge bases that gather published data from preclinical experiments and real-life clinical data are used to assess the potential impact of identified alterations on protein function. However, it is impossible to keep up with the amount of data generated with high-throughput technologies, and many variants lack the functional annotation necessary to distinguish oncogenic drivers from passenger events with little to no significant diagnostic, prognostic, or predictive impact. While the effect of some types of genetic variants, such as frameshift and nonsense variants, is quite definite, it is particularly challenging to predict the outcome of missense variants. This issue was soon recognized and resulted in the development of several algorithms that mainly employ information about evolutionary conservation and sequential or physicochemical properties, which might prove helpful for Mendelian disorders [3]. However, for cancer-associated proteins, a robust prediction requires a more comprehensive assessment that also employs structural data or binding properties of known inhibitors to reliably sort variants that should be pursued in preclinical studies or even clinical scenarios.

## Minimal information for job submission

Computational pipelines that handle biological data can string together multiple software tools, each with its own settings and parameters. This can add multiple layers of complexity barring non-specialist users with little background in bioinformatics to access such a pipeline. Thus, it is important for a clinically relevant tool to have a low barrier to access. Making the tool available as an online web server would make access easier. Therefore, we have developed the new web server PredictONCO, which can become a valuable tool for routine analysis of the data from next-generation sequencing experiments. The tool is designed keeping in view the urgency of oncologically relevant analysis, hence it was made fully automated, with very little input required from a user’s side. Effectively, the only two pieces of information required to start a job on the web server are the target protein’s name and the associated mutation. Inputting this information is done via the easy-to-use graphical user interface of the web server (**Figure 1**). Once the job has started, it can take about a day to complete, but it can be longer with a load of the server. However, if the calculation for that protein and mutation combination has been made previously, the results are provided immediately from the jobs database. All completed calculations are added to the results page as soon as they are available, regardless of the status of the other calculations. An email alert is also sent to the user upon initiation and completion of the calculations, providing identification of the calculation and the hyperlink to the web page with results.

**Figure 1.**
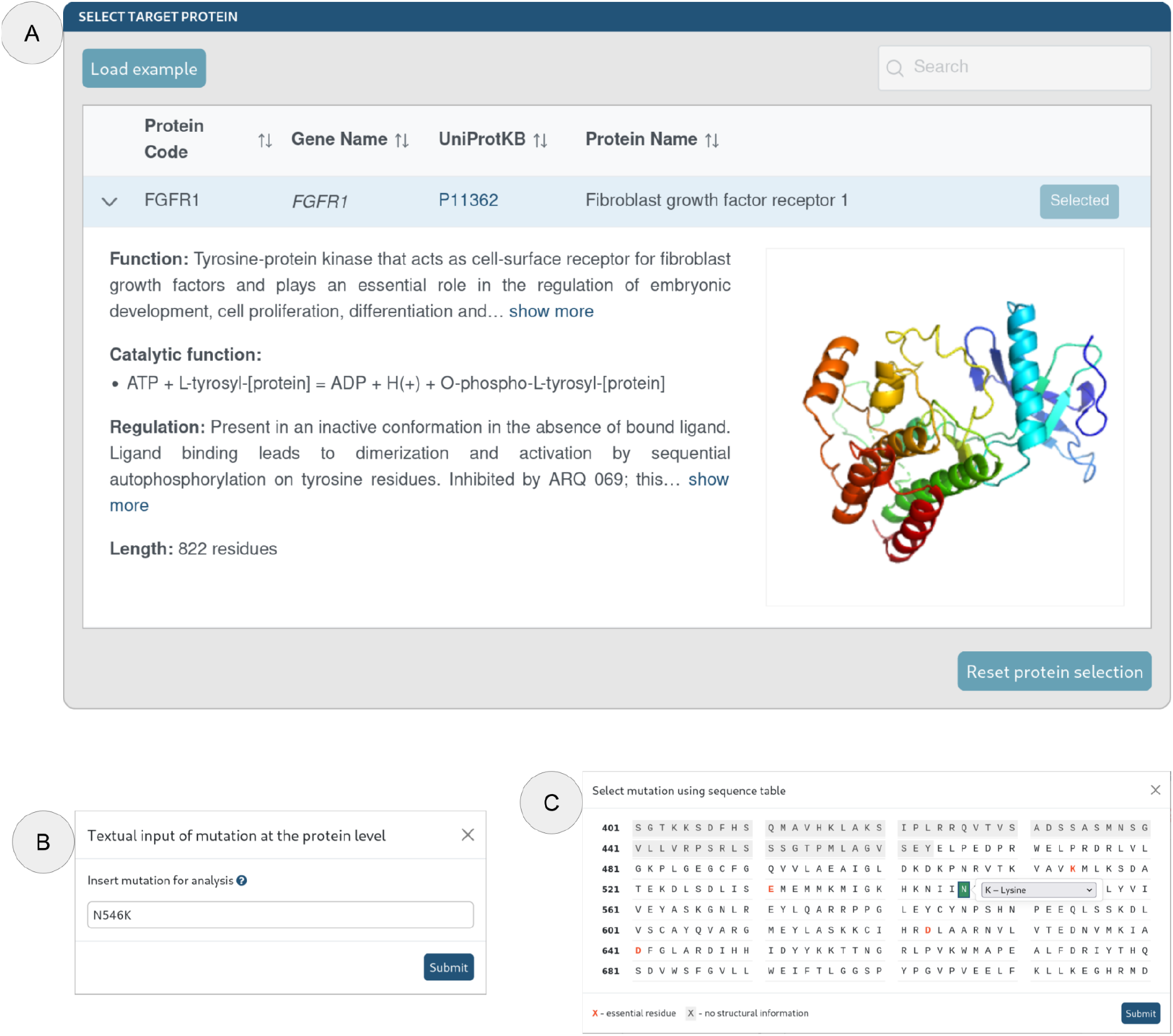
The graphical user interface of PredictONCO web server’s job submission page. (**A**) Protein selection window, (**B**) Mutation selection window via textual input, and (**C**) Alternative mutation selection window via the selection on the sequence table.

## Output information and interpretation of results

The results page is an easy to use collection of calculations, organised in separate fields (**Figure 2**). The wild type structures are used by the pipeline to calculate the stability changes using FoldX [4] and Rosetta [5]. The pipeline also models structures of the mutant proteins, and these modelled structures have pocket level information calculated using P2Rank [6] as well as information about essential residues from M-CSA [7] and UniProt databases [8]. The pKa values of ionizable groups, indicative of changes in reactivity between the wild type and mutant proteins are calculated using PropKa [9]. The newly developed XGBoost-based predictor uses all obtained values as features to return the probability of a mutation’s oncogenic effect. To create the predictor, we used a dataset of 464 oncogenic and 449 non-oncogenic mutations compiled from the ClinVar [10] and OncoKB [11] databases. All mutations were annotated with a clinically verified effect based on available information from precision oncology databases [12-15] and primary literature. The predictor was validated on an independent subset of held-out data, demonstrating areas under the receiver operating characteristic curve (ROC AUC) of 0.95 and 0.94 for structure-based and sequence-based predictions, respectively. The average precision from the Precision-Recall curve was 0.98 and 0.94 for structure-based and sequence-based predictions, respectively.

**Figure 2.**
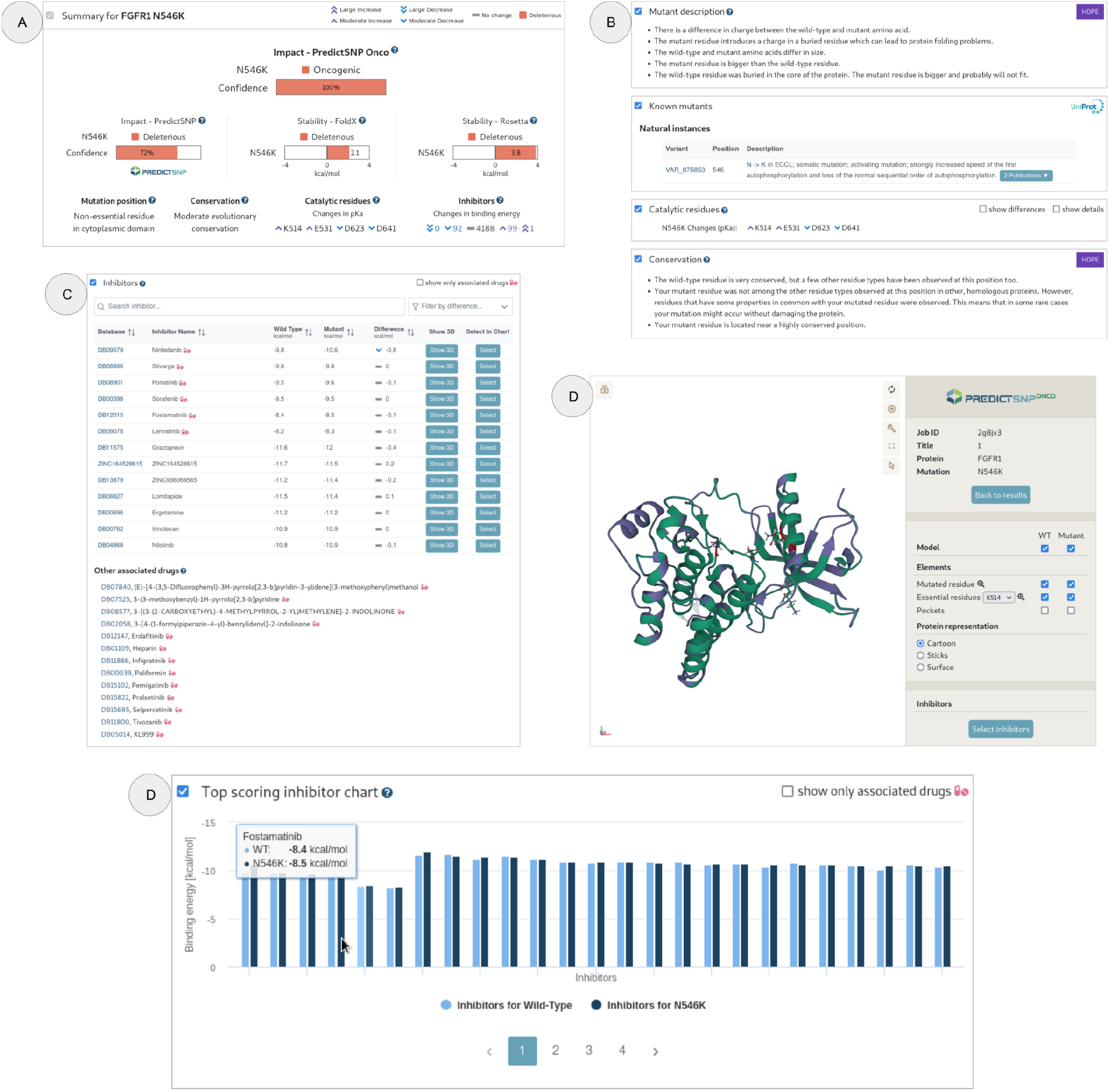
The graphical user interface of PredictONCO web server’s results page. (**A**) An ‘at a glance’ style Summary window, which compiles the most important calculations, (**B**) Various other analyses detailed in their own windows, such as Mutant description, Known mutants, and Conservation analysis from the HOPE server, as well as a window reporting changes in pKa for the catalytic residue, **(C)** Inhibitors window for showing binding energies of inhibitors in the wild type and the mutant protein, (**D**) Protein structure visualisation window for viewing the wild type and mutant protein structures with various settings. This window also allows for the visualisation of inhibitors and other protein features such as pockets and essential residues, and (**E**) The Top scoring inhibitor chart which compares the top 100 binding energies for individual inhibitors as a bar chart.

The results page also contains docking results of 4380 small molecules approved by the Food and Drug Administration and European Medicines Agency, docked onto both the wild type and the mutant structure using Autodock Vina [16]. Changes in the binding affinity of the drugs associated with the target protein upon mutation can support decisions on treatments (**Figure 2**). The structure visualisation page allows users to inspect the tertiary structure of the wild type and mutant protein, along with the mutated residue, essential residues, and bound top-scoring drugs, in various visualisation formats. Furthermore, the results page contains information from other useful tools and databases such as UniProt [8] and HOPE [17], as well as pathogenicity scores based on the PredictSNP server [18]. The most important bits from the results page are available at the top in the ‘Summary’ field, along with the PredictONCO oncogenicity score. This score utilises multiple outputs of the pipeline to predict the mutation’s result on the target protein’s oncogenicity in a single value. To demonstrate the tool’s usage, results for variants in two cancer-associated proteins, cellular tumour antigen p53, and fibroblast growth factor receptor FGFR1, are presented as case studies.

## Case study with R175H and K139M variants of cellular tumour antigen p53

For p53, the R175H and K139M variants were submitted for analysis, with R175H being a notoriously known inactivating variant and K139M being an unknown alteration identified by comprehensive genomic profiling. Input data consisted only of the respective protein selection and selection of a particular mutation through either textual or sequence entry using a nomenclature corresponding with the canonical transcript. For R175H, the PredictONCO results showed a deleterious prediction on both the stability level and by the PredictSNP consensus classifier. Taking all calculations into account, the variant is predicted to be deleterious with a 100% confidence score, which is in agreement with the variant being a well-known cancer-associated event leading to a loss of protein function. Its occurrence in many tumour types, of both germline and somatic origin, is also shown in the “Known mutants” section, which makes the data interpretation-related literature search easier by providing the user with relevant references.

The K139M variant of cellular tumour antigen p53, on the other hand, is a variant that lacks proper functional characterization and requires careful assessment for subsequent clinical management, especially when of germline origin, which makes it suitable for PredictONCO evaluation. PredictSNP consensus classifier predicted a deleterious effect with a moderate confidence score of 61%, while both stability predictors predicted a neutral effect. However, differences in physicochemical properties and reported occurrence of different mutants in identical residues suggest a functional impact. The overall prediction indicates a deleterious effect with a 98% confidence score. Therefore, by not relying just on the results of widely used sequence-based prediction algorithms, we were able to significantly increase the confidence in protein effect prediction, by 37 p.p. specifically. With such increased confidence, the clinical management of patients harbouring this germline variant would support further studies of incidence in the family and potential co-segregation with the disease.

## Case study for N546K variant of fibroblast growth factor receptor FGFR1

A similar example can be applied to known protooncogenes with the added benefit of inhibitor binding data. Demonstrated by the example of the FGFR1 N546K variant, we got an overall deleterious prediction with a 100% confidence score. Similar to the p53 R175H mutant, several literature references showing causality in cancers and an activating effect on protein function were available. Most importantly, as multiple inhibitors (e.g., Nintedanib, Stivarga, Ponatinib, etc.) can target FGFR1, their comprehensive overview was provided. Inhibitors were accompanied by calculated changes in binding energies, whose decreased values suggest better binding capability, which can help in the selection process if multiple options can be considered. All calculations were performed in a reasonably timely manner, under 2 hours for p53 and 6 hours for FGFR1, with the difference being explained by inhibitor docking and binding energy calculations.

## Conclusions

PredictONCO is a web-based tool that uses computational algorithms to predict the effect of somatic alterations in cancer-associated proteins. It employs several algorithms and database searches that assess the impact of a mutation on protein stability, functionality, and oncogenicity. Importantly, PredictONCO also quantifies the effect of mutations on protein-drug interactions. The input for the web server is straightforward, with only the name of the target protein and associated mutation required. The results page contains several fields with different calculated properties of the mutant protein, including structure, stability change, pocket level information, and essential residues. The web server is fully automated, with email alerts sent to users upon initiation and completion of calculations, and all completed calculations are added to the results page as soon as they become available.

## Web tool availability

The service PredictONCO is available free of charge to all users at the website https://loschmidt.chemi.muni.cz/predictonco/.

## Acknowledgements

The authors would like to express their thanks for the Czech Ministry of Education (CZECRIN LM2023049, TEAMING CZ.02.1.01/0.0/0.0/17_043/0009632), the Ministry of Health (NU20-03-00240) the Technology Agency of the Czech Republic (PerMed TN02000109), European Union (TEAMING 857560, NPO EXCELES LX22NPO5102), and Brno University of Technology (FIT-S-23-8209) for financial support.

